# Endogenous Regulator of G protein Signaling 14 (RGS14) suppresses cocaine-induced emotionally motivated behaviors in female mice

**DOI:** 10.1101/2024.09.12.612719

**Authors:** Sara N. Bramlett, Stephanie L. Foster, David Weinshenker, John R. Hepler

## Abstract

Addictive drugs hijack the neuronal mechanisms of learning and memory in motivation and emotion processing circuits to reinforce their own use. Regulator of G-protein Signaling 14 (RGS14) is a natural suppressor of post-synaptic plasticity underlying learning and memory in the hippocampus. The present study used immunofluorescence and RGS14 knockout mice to assess the role of RGS14 in behavioral plasticity and reward learning induced by chronic cocaine in emotional-motivational circuits. We report that RGS14 is strongly expressed in discrete regions of the ventral striatum and extended amygdala in wild-type mice, and is co-expressed with D1 and D2 dopamine receptors in neurons of the nucleus accumbens (NAc). Of note, we found that RGS14 is upregulated in the NAc in mice with chronic cocaine history following acute cocaine treatment. We found significantly increased cocaine-induced locomotor sensitization, as well as enhanced conditioned place preference and conditioned locomotor activity in RGS14-deficient mice compared to wild-type littermates. Together, these findings suggest that endogenous RGS14 suppresses cocaine-induced plasticity in emotional-motivational circuits, implicating RGS14 as a protective agent against the maladaptive neuroplastic changes that occur during addiction.

## 1. INTRODUCTION

Addiction is a disease in which compulsive drug use gives rise to compounding harmful consequences to health and well-being. All known addictive drugs directly or indirectly enhance dopamine signaling in the nucleus accumbens (NAc), a critical nexus for motivated behavior located in the ventral striatum [1,2]. Compulsive intake of addictive drugs is driven at first by the appetitive effects of drug reward, then by the aversive effects of drug withdrawal [3,4]. This involves a shift in processing from the basolateral amygdala (BLA) to subregions of the extended amygdala, including the central amygdala nucleus (CeA) and the bed nucleus of the stria terminalis (BNST), extended amygdala components. The extended amygdala is critically involved in orchestrating responses to negative emotional information, including mediating fear and anxiety-related behaviors, in part due to its extensive interconnections with loci in the hypothalamus and brainstem that control autonomic, neuroendocrine, and behavioral responses to stress [5–8].

Addictive drugs reconfigure the brain in part by engaging mechanisms of synaptic plasticity, including long-term potentiation (LTP) and long-term depression (LTD) of synaptic strength in basal ganglia and limbic regions [9,10,11]. The highly addictive psychostimulant cocaine enhances extracellular levels of dopamine and other monoamine neurotransmitters throughout the brain by directly blocking their presynaptic reuptake to alter both acute neuron excitability and neuroplasticity [12,13,14]. Monoamine neurotransmission in emotional-motivational brain circuits is a major mechanism of adaptive behavior that cocaine usurps to promote its continued use [15–19]. Increased dopamine neurotransmission is especially impactful in emotional-motivational processing regions due to high dopamine receptor expression [20]. Acute behavioral effects of cocaine occur mostly through transient changes in ion channel conductivity, while effects on learning and memory are tied to convergent signaling with glutamate receptors onto activation of phosphorylated extracellular signaling regulated kinase (ERK) [21–24].

Regulator of G-protein Signaling 14 (RGS14) is a member of the RGS family of proteins [25]. In addition to an RGS domain which confers inactivating GTPase-accelerating protein (GAP) activity on GTP-bound Gαi/o proteins, RGS14 also contains tandem binding domains for Ras/Rap (R1 and R2) that bind and sequester active H-Ras and Ca^2+^/calmodulin to suppresses calcium and ERK signaling [26–29]. Studies show that in hippocampal area CA2, RGS14 suppresses post-synaptic ERK-dependent LTP and is also essential for mGluR-dependent LTD [30,31]. Transient increases in ERK activity lead to short-lived enhancements of neuronal excitability mostly through actions at ion channels in dendrites and spines [32,33], while persistent ERK overactivation promotes nuclear translocation and stabilization of neuroplastic changes via modification of transcription factor activity [34,35]. By altering ERK and calcium signaling in CA2 and other regions, evidence suggest that RGS14 has the overall metaplastic effect of suppressing post-synaptic spine growth [27].

Evidence supports the notion that RGS14 suppresses postsynaptic growth related to emotion-driven learning, a critical mechanism of adaptive behavior [36]. A recent study shows enriched RGS14 protein in limbic regions specifically tied to motivation including the NAc, dorsolateral BNST, and CeA [37]. We previously reported that genetic deletion of RGS14 (RGS14-KO) enhances Morris water maze performance in mice [30], while another group reported elevated fear conditioning in RGS14-KO females [38], suggesting that RGS14 loss improves hippocampal- and amygdala-based learning and memory, the latter in a sex-dependent manner.

RGS14-KO mice also exhibit greater locomotor activation and increased anxiety-like behavior in response to combined novelty and acute cocaine, accompanied by higher pERK1/2 and c-Fos immunoreactivity in the hippocampus and CeA [39]. As mentioned above, addictive drugs strongly induce pERK in NAc, CeA, and dorsolateral BNST [40], regions that also highly express RGS14 [37]. Together, these studies suggest that RGS14 may protect against developing addiction by serving as a natural suppressor of synaptic plasticity in emotion and motivation processing circuits, and that RGS14 dysfunction could promote vulnerability to addiction.

To investigate our hypotheses, we examined cocaine-induced behavioral plasticity and reward-based learning in RGS14-KO mice using locomotor sensitization and conditioned place preference (CPP) paradigms. We also assessed RGS14 immunoreactivity in the striatum and extended amygdala in WT mice using confocal microscopy, including examining colocalization with fluorophore-tagged dopamine receptors in the striatum and evaluating signal intensity following cocaine challenge. We focused on female mice in our investigation, according to preliminary evidence that RGS14-KO had more profound effects on cocaine-induced plasticity in females than in males.

## 2. METHODS

### 2.1. Subjects

All animal housing and procedures were designed according to the National Institutes of Health Guidelines for the Care and Use of Laboratory Animals and were approved by the Emory University Institutional Animal Care and Use Committee. Mice homozygous for a RGS14-deleting mutation (RGS14-KO) were generated as described in Lee at al., 2010 and maintained on a C57BL/6J background. Adult (aged 3-7 months) RGS14-KO mice and wild-type (WT) littermates of both sexes were included in preliminary experiments, the results of which guided the decision to include only adult females in the experiments described in this text. All testing groups were comprised of an even distribution of WT and RGS14-KO subjects, as well as *Drd1a*-tdTomato and *Drd2*-EGFP mice (described in Shuen et al. [41] and Gong et al. [42]). Mice were maintained on a 12-hour light/12-hour dark cycle while socially housed with free access to food and water, except during behavioral testing. All subjects were acclimated to housing in the testing room for >1 week prior to beginning experiments.

### 2.2. Drugs

Cocaine hydrochloride (5 mg/kg, i.p.; National Institute on Drug Abuse Drug Supply Program, Bethesda, MD) dissolved in saline, or saline alone, was administered at a volume of 10 mL/kg body weight. Testing began on subjects immediately following injection.

### 2.3. Locomotor sensitization

Subjects (n = 48) were handled for 1-2 min daily for 4 days preceding the start of the experiment. Locomotor chambers consisted of an enclosed 10”×18”×10” transparent polycarbonate box with corn cob bedding thinly layered over the floor. Locomotor chambers were surrounded by an apparatus projecting a horizontal 4×8 infrared photobeam grid connected to a monitoring system that records photobeam interference (Photobeam Activity System, San Diego Instruments, San Diego, CA). Two consecutive beam breaks constituted an ambulation, which were recorded in 5-min bins and used to quantify locomotor activity.

The paradigm consisted of the following consecutive phases: 3 days of habituation, 5 days of induction, 10 days of withdrawal, and 1 day of cocaine challenge. During habituation, all subjects received saline injections and then locomotor activity was recorded for 30 min. During induction, locomotor activity was recorded while subjects habituated to chambers for 90 min, then half of the subjects (12 WT, 12 RGS14-KO) received cocaine injections while the other half received saline and locomotor activity was recorded for an additional 60 min. During withdrawal, subjects were left undisturbed in their home cages. During challenge, locomotor activity was recorded while subjects habituated to chambers for 90 min, then all subjects received cocaine injections and locomotor activity was recorded for an additional 60 min.

### 2.4. Conditioned place preference

Subjects (n = 32) were handled for 1-2 min daily for 7 days preceding the start of the experiment and habituated to injections with saline on the last 3 of the 7 days. Conditioned place preference apparatuses (San Diego Instruments, San Diego, CA) consisted of a 27”×8.375”×13.5” 3-compartment acrylic enclosure with side compartments each of length 10.75” and a center compartment of length 5.5”, separated by removable panels. Side compartments were distinguishable by visual and tactile cues: one side featured horizontally-striped outer walls and a smooth white floor, while the other side featured vertically-striped outer walls and a rough black floor. All surfaces were opaque except for chamber lids, which were transparent to allow for video tracking by overhead cameras. Mouse activity was recorded and processed into compartments defined manually prior to testing using ANY-maze automatic video tracking software (Stoelting Co., Wood Dale, IL). Time and distance in each compartment were used as metrics for place preference and locomotor activity, respectively.

Baseline place preference was established during a pre-test session, in which chamber separation panels were removed and subjects were placed in the center compartment and permitted to freely explore both sides of the apparatus for 15 min. A biased design was chosen for the paradigm, meaning that the side on which a subject spent the least amount of time during pre-test (i.e. the non-preferred side) was paired with cocaine during conditioning and the preferred side was paired with saline. Control subjects received saline on both sides.

Conditioning was carried out on a schedule of 2 sessions per subject per day, spaced ∼4 hours apart, over 3 consecutive days. To avoid lingering psychoactive effects of drug during same-day conditioning sessions, saline pairing was done during the AM session and cocaine pairing was done during the PM session each day. During conditioning, chamber separation panels were put in place and subjects were injected with either cocaine or saline solution, then immediately placed in the side compartment assigned to the given treatment, where they remained for 30 min before being returned to their home cages. Post-conditioning tests (i.e. post-tests) were conducted the day after the final conditioning sessions and at two 10-day intervals thereafter.

Subjects were left undisturbed in their home cages during the periods between post-tests. Post-test procedure was identical to pre-test.

### 2.5. Immunohistochemistry

Tissue was harvested from mice in locomotor sensitization cohorts 2 and 3 (n = 32) either 2 hours or 24 hours following cocaine challenge. Subjects were injected with 200 mg/kg sodium pentobarbital and transcardially perfused with 4% paraformaldehyde in 0.01 M PBS. Extracted brains were submerged in paraformaldehyde solution for 24 hours for post-fixation, then floated in 15% sucrose/PBS for 24 hours followed by 30% sucrose/PBS for ≥24 hours until sunk. Brains were embedded in OCT on dry ice and sliced coronally into 40 µm sections using a cryostat at -20°C. Sections were stored in PBS+0.025% sodium azide at 4°C.

Prior to staining, tissue sections underwent antigen retrieval in 10 mM sodium citrate buffer for 3 min at 100°C. Sections were washed in PBS and blocked for 1 hour in 5% normal goat serum/0.1% Triton-X/PBS, then incubated in mouse α-RGS14 (NeuroMab, Davis, CA, clone N133/21; 1:500) primary antibody in 5% normal goat serum/0.1% Triton-X/PBS at 4°C overnight. Sections were washed again in PBS and incubated in goat α-mouse AlexaFluor 488+ (Invitrogen, 1:500) secondary antibody in 5% normal goat serum/0.1% Triton-X/PBS for 2 hours, washed a final time in PBS, and mounted onto Superfrost Plus slides (Thermo Fisher Scientific, Waltham, MA). Once dry, slides were coverslipped with Fluoromount-G+DAPI mounting medium (Southern Biotech, Birmingham, AL) and stored at 4°C.

For visualization of RGS14 colocalization with D1 and D2 dopamine receptors in the striatum, free-floating paraformaldehyde-fixed tissue sections from *Drd1a*-tdTomato and *Drd2-* EGFP mice were incubated with mouse α-RGS14 primary antibody followed by either goat α-mouse AlexaFluor 488 (for *Drd1-tdTomato* tissue) or goat α-mouse AlexaFluor 568 (for *Drd2-EGFP* tissue) secondary antibody and mounted as described above.

Slides were imaged using a Leica SP8 MP confocal microscope with tiling to stitch together adjacent scans (10% overlap). ImageJ was used for all microscopy image post-processing. The processing pipeline included: despeckle → median filter → remove outliers (for artifact elimination) → adjust brightness/contrast → apply LUT. Occasionally to provide anatomical clarity, processed images were overlaid onto grayscale raw images. Raw files of all images are available upon request. ImageJ was also used to quantify RGS14 by calculating the mean fluorescent intensity within manually drawn regions of interest. Background fluorescent intensity for each image was measured in the medial septal nucleus, which does not contain RGS14, and subtracted from the mean fluorescent intensity of the region of interest. Figures were composed using Inkscape.

### 2.6. Statistics

Statistical tests used to analyze locomotor sensitization and conditioned place preference data are listed in Supplementary Tables 1 & 2. Data was organized and analyzed in R using base R and the following packages: dplyr, tidyr, stats, rstatix, lmer, emmeans, afex, Rmisc, DescTools, outliers. Plots were generated using ggplot2. Tables were generated using Microsoft Excel. Homogeneity of variance was assessed using Levene’s test, normality was assessed using the Shapiro-Wilk test and analysis of residuals, and outliers were assessed using Grubbs’ test. Assumption violations were addressed by rank transformation of data and/or removal of outliers. Given the robustness of linear mixed-effects models to violations of homoskedasticity and normality [43], modest failures of tests for these assumptions were discounted if the model was otherwise a good fit for the data based on residual analysis. Effects testing of linear mixed-effects models used the Satterthwaite method. Post-hoc contrasts were analyzed using estimated marginal means (emmeans package) and the Kenward-Roger approximation was used for contrasts in linear mixed-effects models. For repeated measures analysis of variance (ANOVA), the Greenhouse–Geisser correction was applied in cases of sphericity violations.

Treatment group differences in RGS14 levels in NAc in WT subjects as measured by mean fluorescent density were analyzed by Welch’s t-tests. The alpha level for all tests was 0.05.

## 3. RESULTS

### 3.1. RGS14 is enriched in distinct subregions of the nucleus accumbens, bed nucleus of stria terminalis, and amygdala

We previously identified RGS14 protein in murine striatum and extended amygdala [37,39].

To obtain a more detailed profile of subregion-specific expression, we immunostained for RGS14 in naive WT mice. Tile-stitched fluorescence micrographs were obtained of full brain regions of interest on the coronal plane. We observed high RGS14 immunoreactivity in the NAc core just ventral to the anterior commissure, with more moderate signal seen dorsal to the commissure in a pattern that was essentially continuous with ventromedial caudoputamen (CPu). In contrast, we saw relatively low signal in a patchy distribution within NAc shell (Fig.1A). In the BNST, high signal was seen exclusively in the dorsolateral region in a tightly restricted pattern that closely aligned with reference atlas delineations of the oval and juxtacapsular subnuclei (Fig.1B). In the amygdala, abundant signal was restricted to the lateral CeA (CeL), surrounded by sparse signal in the adjacent areas of the medial and capsular CeA (CeM and CeC). Low signal was also seen in the basolateral amygdala, a subregion involved in assigning positive emotional valence (Fig.1C).

**Figure 1.**
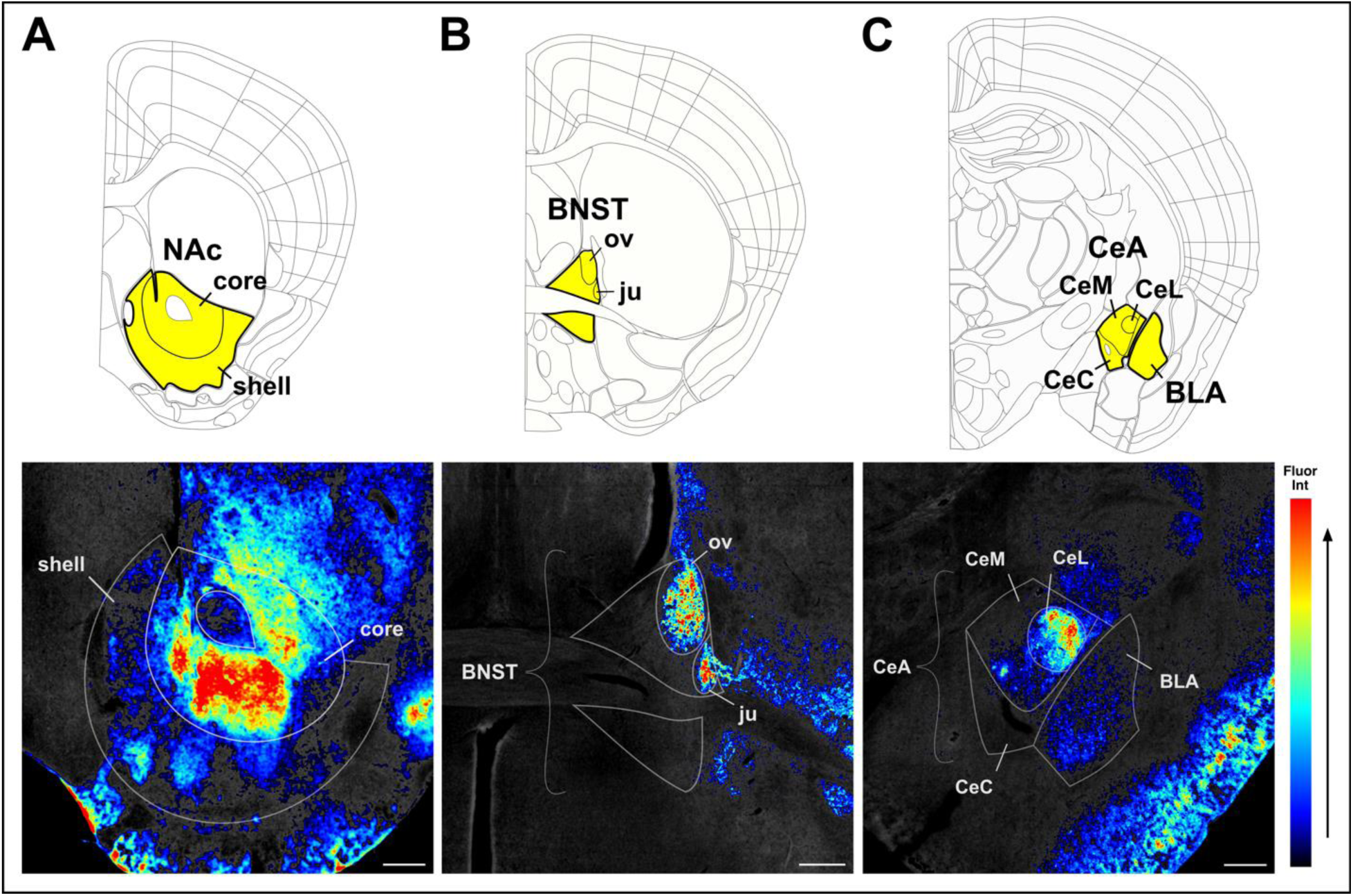
RGS14 is RGS14 is strongly expressed in distinct subregions of the nucleus accumbens, bed nucleus of stria terminalis, and amygdala. Immunofluorescence detection of RGS14 in (A) NAc, (B) BNST, and (C) amygdala in WT mice. Anatomical annotations (top) from the Allen Mouse Brain Reference Atlas (atlas.brain-map.org) aligned with tile-scan images of RGS14-immunostained tissue (bottom), pseudocolored to show relative fluorescence intensity. Highlighted atlas regions are approximately outlined in tissue sections. Sections ordered rostral to caudal. Scale bar = 250 µM. Anatomical abbreviations:: BLA: basolateral amygdala, BNST(ju,ov): bed nucleus of the stria terminalis-(juxtacapsular, oval), CeA: central amygdala, Ce(C,L,M): central amygdala-(capsular, lateral, medial), NAc: nucleus accumbens.

### 3.2. RGS14 is localized to D1 and D2 dopamine receptor-expressing neurons in the nucleus accumbens

Most striatal medium spiny projection neurons exclusively express either D1 or D2 dopamine receptors (D1R & D2R) [44,45]. In the mouse ventral striatum, both D1R- and D2R-containing medium spiny neurons (D1-MSNs & D2-MSNs) project along direct and indirect pathways through the basal ganglia [46], and work in concert in the NAc core to process motivated learning and adaptive behavior [47–51]. As shown in Figure 1A, RGS14 immunoreactivity is abundant in the striatum, particularly within NAc core. Using an RGS14 reporter mouse that targets RGS14 to the nuclei of cells where it is expressed [52], we verified intrinsic RGS14 protein expression in NAc neurons [37]. To assess RGS14 abundance in D1 vs D2 receptor expressing NAc neurons, we immunostained for RGS14 in striatal tissue sections from *Drd1-tdTomato* and *Drd2-EGFP* mice and evaluated colocalization (Fig. 2). There was extensive colocalization of RGS14 with both tdTomato and EGFP in NAc core, and to a lesser extent in NAc shell and ventromedial CPu. RGS14 signal was apparent in both cell bodies and neuropil.

**Figure 2.**
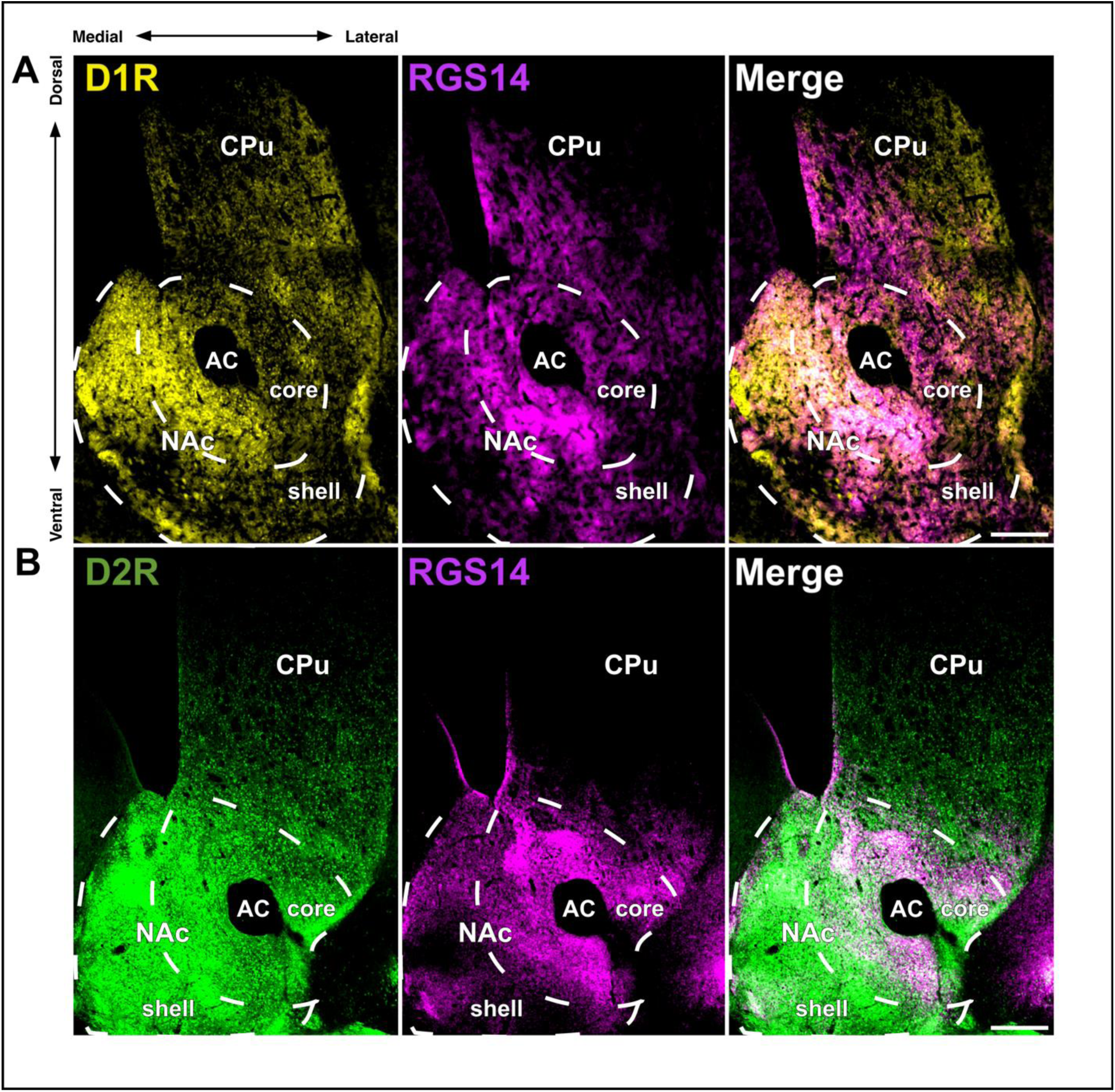
RGS14 colocalizes with D1 and D2 dopamine receptors in nucleus accumbens. Immunofluorescence detection of RGS14 with D1R and D2R in RGS14-immunostained striatal tissue from *Drd1-tdTomato* and *Drd2-EGFP* reporter mice. **A.** *Drd1-tdTomato* tissue immunoreactivity; panels:: <u>yellow (left)</u>: D1R-tdTomato, <u>magenta (middle)</u>: RGS14, <u>white (right</u>): overlap. **B.** *Drd2-EGFP* tissue immunoreactivity:: <u>green (left)</u>: D2R-EGFP, <u>magenta (middle)</u>: RGS14, <u>white (right):</u> overlap. Fluorophores were sequentially scanned in separate channels and computationally merged. Approximate borders of NAc core and shell outlined in white dashes. Scale bar = 250 µM. Anatomical abbreviations:: **CPu**: caudoputamen, **NAc(C,S)**: nucleus accumbens-(core, shell).

### 3.3. RGS14 is upregulated 2 hours after cocaine challenge in the nucleus accumbens of mice with a chronic cocaine history

Our group recently reported that RGS14 is upregulated in the hippocampus by hyperexcitability during status epilepticus [53]. This raises the possibility that RGS14 could also be induced elsewhere to mitigate signaling hyperactivation, such as in the NAc when overactivated by psychostimulants. To determine whether cocaine challenge alters RGS14 induction in mice with a history of cocaine exposure, we quantified RGS14 immunofluorescence intensity 2 h after cocaine challenge (5 mg/kg, i.p.) in subjects 10 days after chronic cocaine administration (5 mg/kg/d, i.p. for 5 days) or saline. We did not find any significant differences in the BNST, CeA, or BLA (data not shown), but RGS14 levels were significantly higher in both the NAc core (*t*(5.50) = 3.48, *p* = .015) and shell (*t*(4.07) = 5.48, *p* = .005) in subjects chronically treated with cocaine compared to saline (Fig. 3).

**Figure 3:**
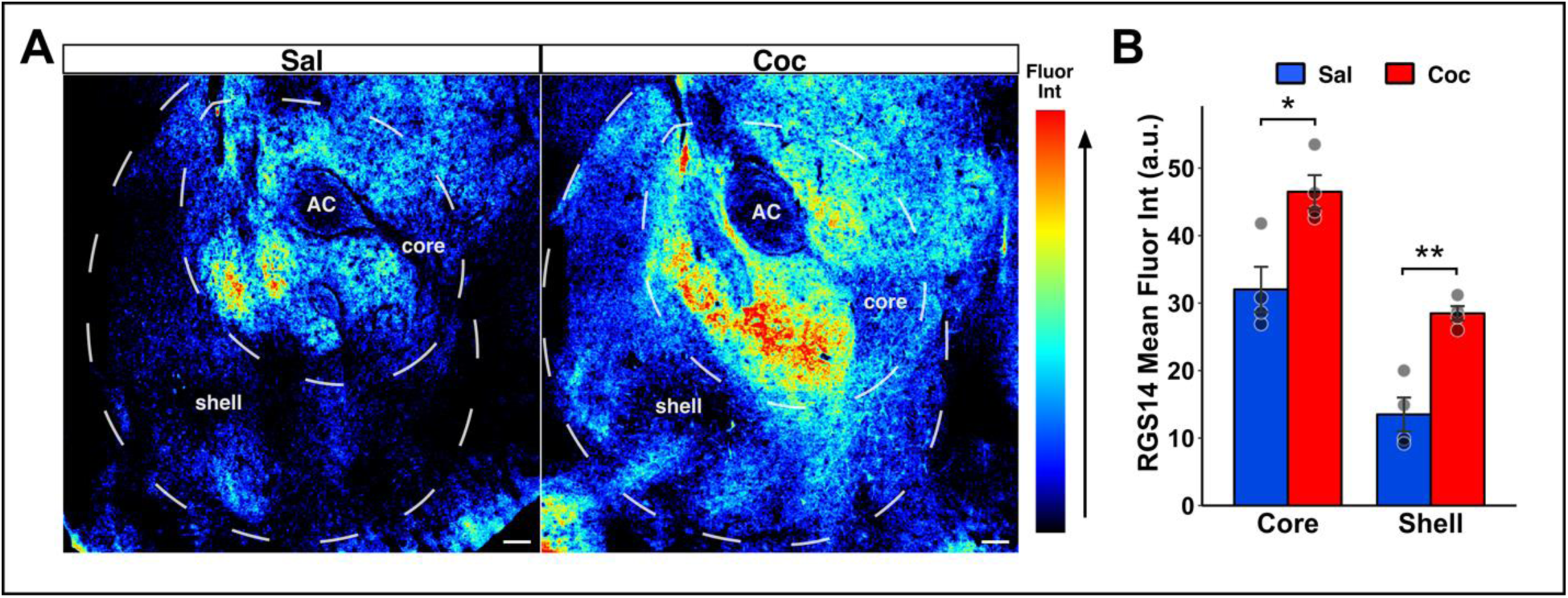
RGS14 is upregulated in nucleus accumbens core and shell 2 hours after cocaine challenge in cocaine-experienced mice. Immunofluorescence detection and relative quantification of RGS14 protein in NAc 2 hours after Coc challenge (i.p.) in naïve and Coc-experienced WT mice. **A.** Representative images of RGS14-immunostained NAc tissue from naive (left) and Coc-experienced (right) mice, pseudocolored to show relative fluorescence intensity. **B.** Mean fluorescence intensity (arbitrary units) in NAc core and shell of RGS14-immunostained tissue. Analyzed by Welch’s t-test. Error bars represent mean ± SEM. Symbols: †p < .10, \**p* < .05, \*\**p* < .01, \*\*\**p* < .001. Scale bar = 100 µm. Anatomical abbreviations:: **AC**: anterior commissure, **NAc(C,S)**: nucleus accumbens- (core, shell).

### 3.4. Enhanced locomotor sensitization to cocaine in RGS14-KO mice

The operational definition of sensitization is a magnitude of response in a subsequent exposure to a stimulus that surpasses that elicited by its initial exposure. It is well-established that locomotor activation sensitizes to repeated cocaine treatment in rodents, an adaptation marked by neuroplastic changes in mesocorticolimbic pathways [54,55] including increased spine density specifically in NAc core [56]. RGS14 inhibits spine enlargement following two-photon glutamate uncaging in CA2 dendrites [27], leading us to suspect that it can also act in the

NAc core, through this and/or additional mechanisms, to suppress cocaine-induced plasticity underlying locomotor sensitization. To test the hypothesis that RGS14 loss increases susceptibility to the behaviorally sensitizing effects of cocaine, we performed the locomotor sensitization experiment diagrammed in Fig. 4A with female RGS14-KO mice and WT littermate controls (n = 48). Briefly, after 3 days of habituation with saline, WT and RGS14-KO mice were treated with either saline or cocaine every day for 5 days (induction of sensitization), withdrawn for 10 days, then given a challenge dose of cocaine (expression of sensitization). Locomotor activity was measured for 60 min following each treatment. We selected a low dose of cocaine (5 mg/kg) (1) to minimize genotype differences in the baseline response to drug in naïve animals, according to our previously published dose response curve [39], and (2) because we predicted that RGS14 KO mice would have enhanced sensitization and wanted to avoid a ceiling effect. For a complete lists of statistical tests, see Tables S1&2.

**Figure 4.**
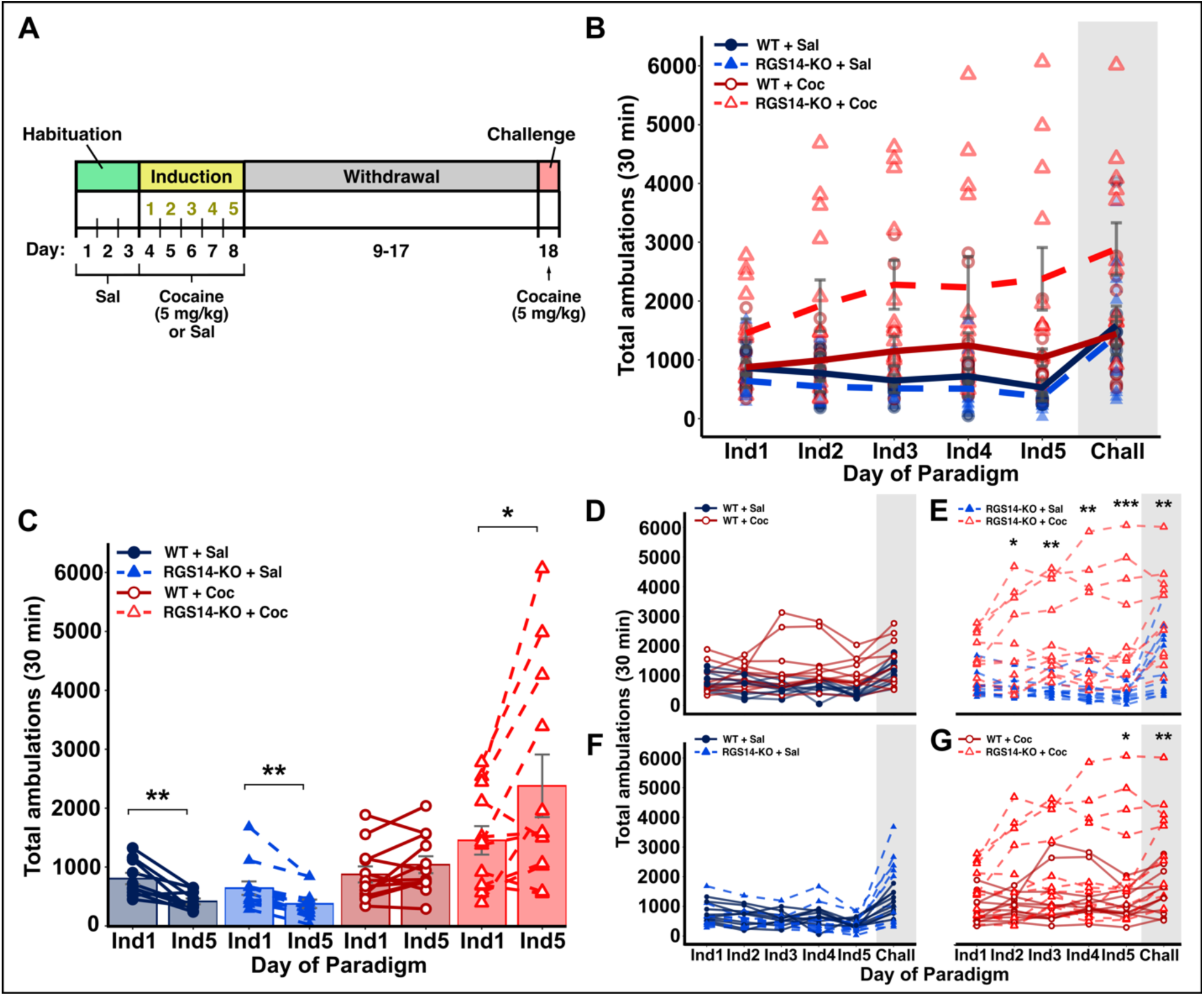
Locomotor sensitization to cocaine is enhanced in RGS14-KO mice. **A.** Diagram of cocaine sensitization protocol (n = 48); cocaine (5mg/kg) or saline administered by i.p. injection. **(B-G)** Total ambulatory activity (beam breaks/30 min) each day of induction and challenge. **B.** Total ambulatory activity (error bars: ± S.E.M.) each day for all groups. There is a significant genotype × treatment × day interaction effect over induction (LMM) and a significant genotype × treatment interaction effect during cocaine challenge (Two-way ANOVA). **C.** Within-subjects changes in ambulatory activity from the first to the last day of induction (Ind1 to Ind5). Locomotor activity was decreased in Sal-treated controls as expected from habituation. Locomotor activity was significantly increased only in the RGS14KO+Coc group (paired pairwise t-tests^).**D.** In WT mice, there is a significant treatment × day interaction effect over induction (LMM), but no significant differences between WT+Coc and WT+Sal group locomotor activity were detected on any individual day (pairwise t-tests^). **E.** In RGS14-KO mice, there is a highly significant treatment × day interaction effect over induction (LMM). Locomotor activity is significantly elevated in the RGS14KO+Coc group over RGS14KO+Sal controls at Ind2-Ind5 (pairwise t-tests^) and challenge (Tukey HSD test). **F.** No significant genotype × day interaction in the Sal treatment group (LMM). **G.** In the cocaine treatment group, there is a significant genotype × day interaction effect over induction (LMM). Locomotor activity is significantly elevated in the RGS14KO+Coc group over WT+Coc controls at Ind5 (pairwise t-tests^) and challenge (Tukey HSD test).

Locomotor activity on each day of the paradigm during the first 30 min following cocaine or saline treatment is shown in Fig. 4B. We first assessed the effects of RGS14 loss on induction of locomotor sensitization to cocaine, and a linear mixed model analysis revealed a significant genotype × treatment × day interaction (*F*(1, 188) = 4.64, *p =* .032). Analysis of locomotor activity on the first and last days of the induction period (Fig. 4C, paired t-tests) showed that chronic cocaine induced significant locomotor sensitization only in RGS14-KO mice (RGS14KO+Coc: *t*(11) = 2.54, *p =* .028); cocaine-induced locomotor activation did not significantly change between the first and final induction phase cocaine treatment in WT controls (WT+Coc: *t*(11) = 1.63, *p =* .131). Saline-treated controls of both genotypes became hypoactive as they habituated, as expected (WT+Sal: *t*(11) = -4.15, *p =* .004; RGS14KO+Sal: *t*(11) = -4.33, *p =* .003), We analyzed group interactions across the induction phase using linear mixed models followed by pairwise comparisons of estimated marginal means. There was a significant treatment × day interaction effect within the WT group (Fig. 4D; *F*(1, 94) = 11.56, *p* < .001), but no significant differences in locomotor activity between cocaine- and saline-treated mice on any individual induction day were detected. We observed a trend for cocaine-induced locomotor activation starting on induction day 3, but it never reached significance (Ind3: *t*(38.1) = 2.34, *p =* .094; Ind4: *t*(38.1) = 2.45, *p =* .094; Ind5: *t*(38.1) = 2.39, *p =* .094). This result is not entirely surprising, as we were using a low dose of cocaine (5 mg/kg) that is below the typical threshold for sensitization in rodents [57,58]. There was a significant treatment × day interaction within the RGS14-KO group (Fig. 4E; *F*(1, 94) = 25.41, *p* < .001). A trend for increased locomotor activity in cocaine-treated mice was evident on induction day 1 (Ind1: *t*(27.8) = 1.79, *p =* .085), and the difference reached statistical significance on induction day 2 (Ind2: *t*(27.8) = 3.03, *p =* .010), after which cocaine consistently elicited significant locomotor activation for the rest of the induction phase (Ind3: *t*(27.8) = 3.88, *p =* .002; Ind4: *t*(27.8) = 3.80, *p =* .002; Ind5: *t*(27.8) = 4.41, *p* < .001).

Within cocaine-treated mice (Fig. 4G), we found a significant genotype × day interaction effect over induction (*F*(1, 94) = 6.75, *p =* .022). Post-hoc analysis confirmed that locomotor activation by acute cocaine was not significantly different between drug-naïve WT and RGS14-KO mice (Ind3: *t*(28.7) = 1.21, *p =* .236), but revealed a trend for enhanced locomotor activation in RGS14-KO mice by the third cocaine exposure (Ind3: *t*(28.7) = 2.37, *p =* .100) that reached statistical significance on the final day of induction (Ind5: *t*(28.7) = 2.80, *p =* .045). Together, these data indicate that loss of RSG14 promotes the induction of sensitization by cocaine.

### 3.5. RGS14 loss increases the expression of cocaine-induced locomotor sensitization

To determine whether RGS14 loss affects the persistent expression of cocaine sensitization, all groups were challenged with cocaine (5 mg/kg, i.p.) 10 d after the last induction day. Locomotor activity was recorded for 60 min. A two-way ANOVA revealed a significant genotype × treatment interaction (*F*(1, 44) = 4.63, *p =* .037) (Fig. 4B, shaded box). Tukey tests revealed that the locomotor response to cocaine challenge was enhanced by chronic cocaine only in RGS14-KO mice (Fig. 4E, shaded box; *t*(44) = 3.41, *p =* .008); locomotor activity in WT mice did not differ by drug history (Fig. 4D, shaded box; *t*(44) = 0.71, *p =* .894). Additionally, the locomotor response was significantly larger in RGS14-KO mice previously given chronic cocaine than in WT mice with the same drug history (Fig. 4F, shaded box; *t*(44) = 3.34, *p =* .009). These data indicate that the augmenting effect of RGS14 loss on the sensitized response to cocaine persists after a period of withdrawal.

### 3.6. RGS14 loss augments the magnitude and duration of cocaine conditioned place preference

We and others showed previously that RGS14 loss enhances spatial learning in mice [30] and fear conditioning specifically in female mice [38]. Based on these studies and the brain expression profile of RGS14 [37], we anticipated that reward learning would be enhanced in female RGS14-KO mice. We tested this hypothesis using a canonical biased cocaine conditioned place preference (CPP) paradigm, except that rather than a single posttest the day after the final conditioning day, we conducted two additional posttests at 10 d intervals thereafter to assess the persistence of the cocaine-association context memory (Figure 5A). We also examined the change in distance traveled on the cocaine-paired side as a measure of conditioned locomotor activation. Refer to Table S3 for full statistical analyses.

**Figure 5.**
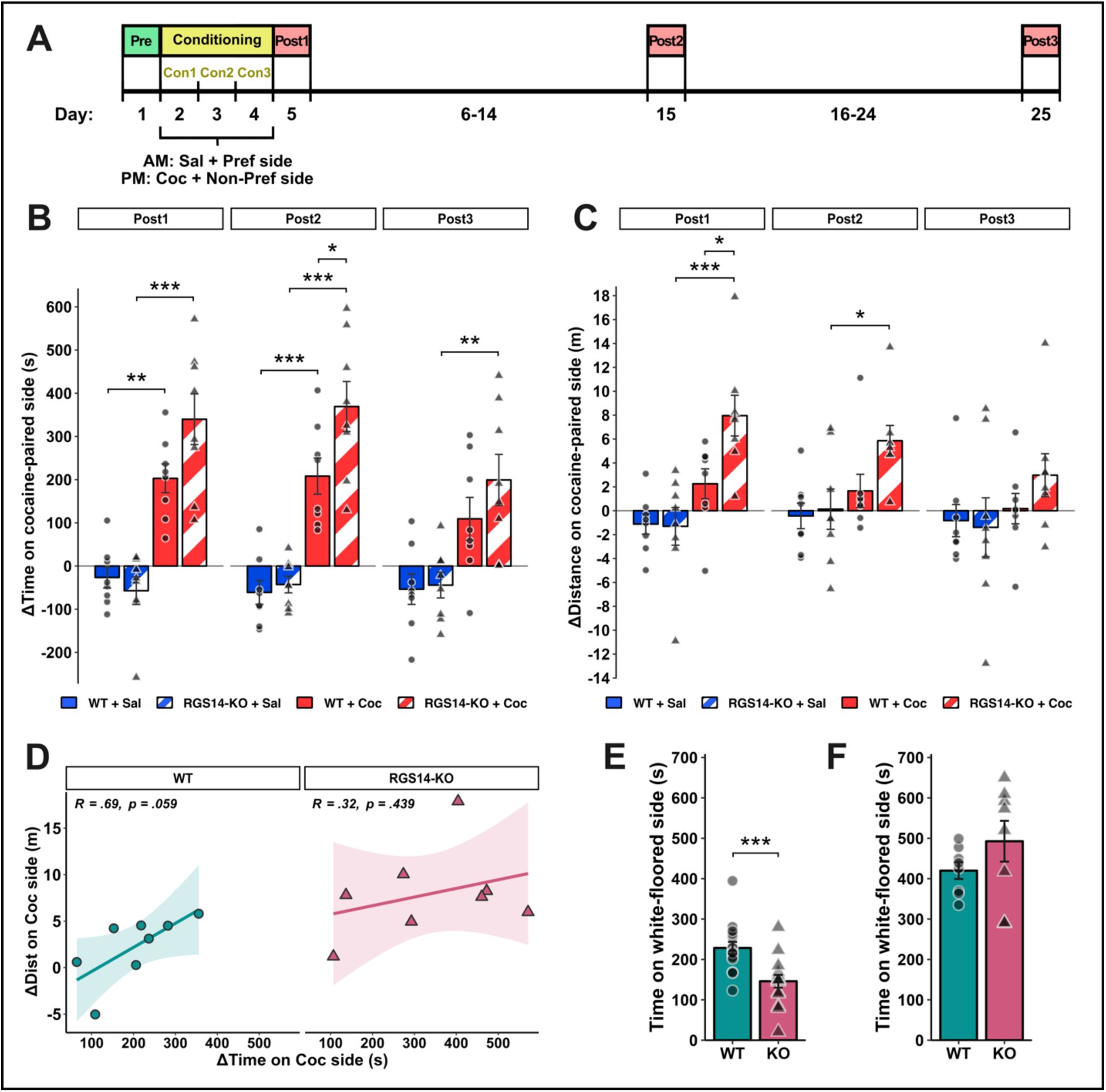
RGS14 loss enhances cocaine-conditioned place preference and locomotor hyperactivity. **A.** Diagram of cocaine-conditioned place preference (CPP) protocol (n = 32); cocaine (5 mg/kg) or saline administered by i.p. injection. **B.** *Increased magnitude and duration of cocaine-conditioned place preference in RGS14-KO mice.* Time spent on the cocaine-paired side (i.e. preference, Δpre-test) was significantly enhanced in WT and RGS14-KO groups at Post1 and Post2, an effect that is significant only in RGS14-KO mice at Post3. Preference was significantly enhanced in RGS14-KO mice over WT controls at Post2 (Tukey HSD test^). **C.** *Increased magnitude and duration of cocaine-conditioned locomotor hyperactivity in RGS14-KO mice.* Distance traveled on the cocaine-paired side (Δpre-test) was significantly enhanced exclusively in RGS14-KO mice at Post1 and Post2. ΔDistance was significantly enhanced in RGS14-KO mice over WT controls at Post1 (Tukey HSD test^). **D.** Correlation plots of cocaine-conditioned place preference (Δtime) and locomotor hyperactivation (Δdistance) at Post1 (shaded area: 95% confidence interval). There is a trend for a positive correlation in WT mice that is absent in RGS14-KO mice (Pearson’s test). **E.** RGS14-KO mice spend significantly less total time in the aversive white-floored compartment during pre-test (n = 32, Welch Two Sample t-test). **F.** *Association with cocaine overcomes stronger inherent place aversion in RGS14-KO mice.* No significant difference between WT and RGS14-KO groups in total time in the white-floored compartment once paired with cocaine (Post1, n = 16, Welch Two Sample t-test). All error bars represent mean ± SEM. Symbols: †p < .10, \**p* < .05, \*\**p* < .01, \*\*\**p* < .001. Carat (^) indicates Bonferroni correction.

CPP results are shown in Figure 5B. When examining all post-tests by three-way repeated measures ANOVA, the main effects for treatment and post-test were significant (Treatment: *F*(1, 28) = 59.24, *p <* .001; Post-test: *F*(1.54, 43.24) = 11.10, *p <* .001), and there was a trend for a main effect of genotype (*F*(1, 28) = 2.99, *p* = .095). The genotype × treatment × post-test interaction effect was not significant (*F*(1.54, 43.24) = 1.24, *p* = .353), but the treatment × post-test interaction was significant (*F*(1.54, 43.24) = 10.78, *p <* .001) and there was a trend for a genotype × treatment interaction (*F*(1, 28) = 3.08, *p* = .090). Two-way ANOVAs at each post-test revealed a significant genotype × treatment interaction effect at Post1 (*F*(1, 28) = 4.57, *p* = .042), a trend for an interaction at Post2 (*F*(1, 28) = 3.26, *p* = .082), and no significant interaction at Post3 (*F*(1, 28) = 0.81, *p* = .377). Main effect of treatment was significant at each post-test (Post1: *F*(1, 28) = 63.88, *p* < .001; Post2: *F*(1, 28) = 74.40, *p* < .001; Post3: *F*(1, 28) = 20.45, *p* < .001).

Analysis of planned comparisons at Post1 revealed that time spent on the cocaine-paired side was significantly enhanced in the cocaine-treated WT (*t*(28) = 4.14, *p* = .002) and RGS14-KO groups (*t*(28) = 7.16, *p* < .001) compared to controls that received saline on both sides, indicating that cocaine induced a place preference in both genotypes. Additionally, there was a trend for a CPP of greater magnitude in RGS14-KO mice (*t*(28) = 2.47, *p* = .087). CPP persisted when measured at Post2 in both WT (*t*(28) = 4.82, *p* < .001) and RGS14-KO groups (*t*(28) = 7.38, *p* < .001), and the initial trend for a greater effect in the RGS14-KO group reached significance at this time point (*t*(28) = 2.88, *p* = .036). By Post3, a full 3 weeks after the final conditioning session, preference for the cocaine-paired side in the WT group had been reduced to a trend (*t*(28) = 2.56, *p* = .072) but remained significant in the RGS14-KO group (Post3: *t*(28) = 3.26, *p* = .004). Together, these data indicate that RGS14 loss increases the magnitude and persistence of cocaine-CPP.

### 3.7. RGS14 loss augments the magnitude and duration of cocaine-conditioned locomotor hyperactivity

Environmental context has been shown to modulate cocaine-induced structural plasticity in NAc core and subsequent psychomotor sensitization [56,59]. Furthermore, cocaine-associated environments can elicit conditioned locomotor hyperactivity in the absence of the acute pharmacological effects of the drug itself [60–63]. Conditioned locomotor hyperactivity may therefore be an indicator of the modulatory effect of a given environment on cocaine-induced plasticity in the NAc and/or NAc-projecting regions. We examined locomotor activity/distance traveled in the cocaine-paired context during CPP post-tests to assess how RGS14 impacts cocaine-conditioned locomotor activation.

Locomotor activity results are shown in Figure 5C. We initially analyzed data from all post-tests using three-way repeated measures ANOVA on ranks. We found no significant genotype × treatment × post-test interaction effect (*F*(1.67, 46.86) = 0.45, *p* = .604), but a trend for a genotype × post-test interaction (*F*(1.67, 46.86) = 3.17, *p* = .060) and a significant main effect of treatment (*F*(1, 28) = 13.11, *p* = .001). Two-way ANOVAs at each post-test showed no significant genotype × treatment interaction effects, but a trend for an interaction at Post1 (*F*(1, 28) = 3.08, *p* = .090). Main effect of treatment was significant at each post-test (Post1: *F*(1, 28) = 29.45, *p* < .001; Post2: *F*(1, 28) = 6.96, *p* = .013; Post3: *F*(1, 28) = 4.46, *p* = .044), and there was a significant main effect of genotype at Post1 (*F*(1, 28) = 5.05, *p* = .033) and a trend at Post2 (*F*(1, 28) = 3.04, *p* = .092).

Analysis of planned comparisons at Post1 revealed that distance traveled on the cocaine-paired side was significantly increased in the cocaine-treated RGS14-KO group (*t*(28) = 5.08, *p* < .001) compared to RGS14-KO controls that received saline on both sides, but there was only a trend for an increase in the cocaine-treated WT group (*t*(28) = 2.60, *p* = .067). Within the cocaine treatment group, locomotor hyperactivity was significantly greater in RGS14-KO mice (*t*(28) = 2.83, *p* = .040) than WTs. At Post2, locomotor activity remained significantly elevated in the cocaine-treated RGS14-KO group compared to saline-treated controls (*t*(28) = 2.74, *p* = .049).

By Post3, no significant differences in locomotor activity were observed between any of the groups. To ensure that observed increases in locomotor activity were specifically elicited by the cocaine-paired context and were not the result of general locomotor hyperactivity, we examined the change in total distance traveled in all compartments and found no significant differences between groups (Fig. S2). Together, these data indicate that RGS14 loss enhances the magnitude and duration of conditioned locomotor hyperactivity in a cocaine-paired context. This suggests that RGS14 suppresses environment-based enhancement of cocaine-induced plasticity in the NAc and/or connected regions.

### 3.8. RGS14 regulates cocaine-conditioned place preference and cocaine-conditioned locomotor activation through dissociable mechanisms

The mechanisms that process emotional valence and behavioral engagement are distinct, but overlap in their governance of motivated behavior. Having determined that RGS14 loss enhances both cocaine-CPP and cocaine-conditioned locomotor hyperactivity, we next asked whether these metaplastic behavioral effects of RGS14 loss were mechanistically intertwined by analyzing the correlation (Pearson’s test) between the change in time and the change in distance on the cocaine-paired side during Post1, when these measures were enhanced in both cocaine-treated WT and RGS14-KO groups (Fig. 5D). We observed a trend for a positive correlation in the WT group (*r*(6) = .69, *p* = .059) but no relationship in the RGS14-KO group (*r*(6) = .32, *p* = .439), suggesting that RGS14 regulates cocaine-CPP and cocaine-conditioned locomotor activation through dissociable mechanisms. This interpretation is further supported by the distinct temporal profiles of cocaine-CPP and cocaine-conditioned locomotor hyperactivity in the RGS14-KO group when compared across post-tests: CPP persists for at least 3 weeks, while hyperactivity decays more rapidly.

### 3.9. Cocaine conditioning overcomes stronger inherent place aversion in RGS14-null mice

While reward is typically associated with inducing positive emotional states, relief from negative emotional states is also rewarding in and of itself; certain anxiolytic drugs like benzodiazepines and the type-1 cannabinoid receptor agonist arachidonylcyclopropylamide support a CPP in rodents [64,65]. The compartments of the CPP chambers used in this study were designed to be emotionally neutral, but one side features a semi-reflective white floor that somewhat augments ambient illumination, potentially making the compartment mildly aversive to rodents [66]. Indeed, nearly all subjects preferred the black-floored compartment at baseline. Based on prior studies showing that RGS14-KO mice are prone to anxiety-like behaviors and may have heightened aversion sensitivity [30,38,39], we predicted that RGS14-KO would enhance baseline aversion to the white-floored compartment in experimental mice. Analysis by Welch’s t-test revealed that RGS14-KO mice spend significantly less time in the white-floored compartment during pre-test than WT controls (Fig. 5E, *t*(28.82) = -3.79, *p* < .001), supporting the notion that RGS14 loss increases sensitivity to aversive stimuli and augments avoidance behavior. Furthermore, time spent in the white-floored compartment post-cocaine pairing was at least as high in the RGS14-KO group as WT (Fig. 5F; *t*(9.28) = 1.33, *p* = .214), indicating that enhanced cocaine-CPP in RGS14-KO mice may be driven, at least in part, by overcoming stronger inherent place aversion.

## 4. DISCUSSION

In the present study, we used behavioral assays and immunohistochemical analyses of genetically modified RGS14-null mice to investigate the role of RGS14 in cocaine-induced behavioral plasticity in emotional-motivational circuits. We observed strong RGS14 immunoreactivity in distinct, subregion-specific patterns in the NAc, dorsolateral BNST, and CeA. We revealed RGS14 immunoreactivity in D1R- and D2R-containing cells in the NAc, where we also found upregulated RGS14 two hours after cocaine challenge in WT mice with a history of chronic cocaine. Lastly, we found that loss of RGS14 augments the magnitude and duration of several behavioral indicators of cocaine-induced psychomotor sensitization and reward learning, including locomotor sensitization, CPP, and conditioned locomotor activation. Together, these findings suggest that RGS14 is a natural and possibly inducible suppressor of cocaine-induced potentiation in emotional-motivational circuits.

### 4.1. RGS14 actions in the nucleus accumbens and cocaine-induced locomotor sensitization

We previously demonstrated RGS14 expression in postsynaptic striatal neurons using an RGS14 nuclear-trapping reporter mouse line [37]. Here, we found overlap of RGS14 immunoreactivity with D1R- and D2R-expressing neurons in the striatum (Fig. 2), enriched in the NAc core, strongly suggesting that RGS14 is present in both direct and indirect pathway MSNs. It has been shown that NAc core D1-MSNs communicate stimulus intensity, while core D2-MSNs signal output prediction error, and they work in concert to process stimulus-outcome associative learning (a.k.a. Pavlovian learning) and control adaptive behavior [51].

Striatal MSNs project through the basal ganglia to drive motor behavior [67]. Cocaine elicits motor hyperactivity by simultaneously stimulating motor-activating D1-MSNs via D1R-Gs/olf and inhibiting motor-inhibiting D2-MSNs via D2R-Gi/o [68]. RGS14-KO mice show increased sensitivity to acute cocaine-induced locomotor activation at a high dose of cocaine (20 mg/kg) [39], presumably because RGS14 acts as a GAP in D2-MSNs to terminate D2R-Gi/o signaling and relieve cocaine-induced suppression of immobility pathways. Unlike in D2-MSNs, it is unlikely that RGS14 modulates cocaine-induced dopamine signaling directly in D1-MSNs due to GAP incompatibility with D1R-Gs/olf. RGS14 is, however, capable of modifying cocaine-induced plasticity in D1-MSNs via binding active H-Ras and/or Ca2+/calmodulin to suppress ERK and/or Ca2+ pathway signaling [26–30]. Response intensity is partially a function of stimulus intensity, which is processed via NAc core D1-MSNs [51]. The locomotor response to the low dose of cocaine used in this study (5 mg/kg) does not significantly differ between drug-naive WT and RGS14-KO mice, but becomes elevated in the RGS14-KO group following repeated cocaine exposure (Fig. 4G). While speculative, the results presented herein hint that RGS14 could be suppressing cocaine-induced potentiation of D1-MSNs in the NAc core to blunt sensitization of the locomotor response. Further study into the role of RGS14 in striatal D1- and D2-MSNs is needed.

### 4.2. RGS14 actions outside of the nucleus accumbens and cocaine-conditioned place preference

We also observed dense RGS14 immunoreactivity in discrete subnuclei of the extended amygdala, namely the oval and juxtacapsular nuclei of the BNST (Fig. 1B), and CeL (Fig. 1C). The extended amygdala connects extensively with brainstem nuclei and the basal ganglia to coordinate physiological and behavioral responses to stress, including evoking the emotional states of fear and anxiety [69,70]. Interestingly, the CeL and BNST oval nucleus specifically contain the largest populations of extrahypothalamic neurons that produce the stress neuropeptide corticotropin-releasing factor (CRF) in the rodent brain [71,72]. Neuropeptides such as CRF can induce significant changes in brain and behavior due to their capacities for prolonged and wide-reaching signaling events [73]. Evidence shows that dysregulation of CRF and other stress systems by chronic and early life stress underlies vulnerability to anxiety and depressive-like symptoms in adulthood [74,75].

We found that drug-naïve RGS14-KO mice were significantly more avoidant of the mildly aversive white-floored CPP compartment (Fig. 5E), suggesting that RGS14 suppresses anxiety-like responses to stressful stimuli. Given that RGS14 inhibits the response to an acute and inherent stressor, RGS14 is likely modifying transient neuronal excitability by negatively modulating receptor-Gi/o signaling. A likely candidate for RGS14 GAP modulation under conditions of stress is CRF type I receptor (CRFR1), which couples with Gi/o when ambient CRF levels are high [76,77]. CRFR1 is widely expressed in the brain, including high local expression in CRF-releasing extended amygdala subnuclei [78,79,80]. In fact, CRFR1 overactivation in the BNST oval nucleus during stress was found to mediate female-specific effects of anxiety-like and avoidance behaviors [80], underscoring the importance of proper CRFR1 regulation in evoking appropriate emotionally-driven responses under stress [81] particularly in females. For this reason and for other reasons discussed below, conditions impairing RGS14 function could be a greater risk factor for developing anxiety disorders in women than men.

Interestingly, we found that association of the white-floored compartment with cocaine reward during conditioning fully abolished stronger avoidance of that compartment by the RGS14-KO group (Fig. 5F). While the exact cell populations and mechanisms driving this shift are unclear, this result has broader implications regarding RGS14 dysfunction creating conditions that precipitate specific vulnerabilities to psychostimulant addiction. Our data shows that RGS14 loss exacerbates inherent avoidant and anxiety-like responses to an aversive context, but also facilitates reversing those responses by permitting stronger association of context with cocaine reward. If these results translate to humans, it suggests that RGS14 dysfunction could underlie a vulnerability to developing psychostimulant addiction through attempted self-medication for anxiety disorders Additionally, increased aversion sensitivity and stress system dysregulation resulting from impaired RGS14 function could heighten vulnerability to the negative affective motivational components of drug withdrawal that drive relapse [3,4].

### 4.3. Dissociation of RGS14 actions on cocaine-induced limbic and motor system adaptations

Another illuminating finding from these data is that RGS14 regulates learning/memory in emotional- and motor-processing circuits in a dissociable manner. Both CPP and conditioned locomotor hyperactivation elicited by cocaine were enhanced by RGS14 loss (Fig. 5B-C), but were not correlated in RGS14-KO mice (Fig. 5D) and decayed on separate time scales. That is, CPP persisted for longer than conditioned locomotor hyperactivation. This suggests that RGS14 effectively inhibits the capacity of cocaine to rewire behavior by suppressing cocaine-induced associative learning based in limbic regions and basal ganglia separately, and that consequences of RGS14 loss on cocaine-induced plasticity in limbic regions controlling emotional associations are likely the most profound. We propose that RGS14 is working in separate populations of neurons to prevent hijacking of the emotional and motor components of goal-directed behavior by cocaine, and is especially critical for regulating emotional drive.

### 4.4. Relevance to human disease: addiction and anxiety in women and girls

The present study points to several intriguing implications for RGS14 in addiction and anxiety disorder pathology, particularly in females. Previous studies demonstrated that loss of RGS14 increased fearful and anxiety-like behaviors in a female-specific manner [38]. Our studies here focused on females, and our findings suggest that RGS14 dysfunction may increase susceptibility to cocaine-induced plasticity in females. The female RGS14-KO mice used in this study demonstrated exaggerated avoidance of the mildly aversive white-floored CPP compartment during pre-test, but pairing the compartment with cocaine reward reversed place preference. Anxiety disorders are more prevalent in women [82], and it has been reported that women are more likely to consume drugs of abuse to self-medicate for anxiety [83,84,85].

Additionally, women are more likely to report stress and negative emotions prior to relapse [86]. RGS14 dysfunction may exacerbate female-specific hypersensitivity to negatively reinforcing stimuli, compounding vulnerability to self-medicating and relapse. Further investigation into the sex-specific effects of RGS14 in addiction- and anxiety-related behaviors is required. Regardless of sex or negativity bias, the results presented herein demonstrate that RGS14 loss is permissive of cocaine-induced behavioral plasticity, suggesting that RGS14 may serve as a protective factor against psychostimulant addiction.

### 4.5. Caveats and limitations

Several of our studies here have limitations that open up opportunities for future exploration. For example, our findings of behavioral sensitization were used as a proxy for plasticity in emotional-motivational circuits. While behavioral sensitization is typically a reliable indicator of synaptic plasticity [87] and we have directly shown that RGS14 loss increases LTP in hippocampal area CA2 [30], synaptic strength itself could be measured in future studies to rule out other mechanisms at play. Locomotor sensitization is commonly used as a reliable measure of psychomotor sensitization though other psychomotor behaviors also sensitize as well (e.g. head movements, stereotypy) [88,89,90] and these could be explored to expand this story. Since these studies were performed with global RGS14 knock outs, other factors could be contributing to the behaviors such as altered basal ganglia function, or RGS14 actions in the periphery such as in the kidney and intestines [91] altering cocaine metabolism, and/or possibly additional mechanisms. Targeted knock-out of RGS14 in brain sub-regions would be informative. Based on preliminary studies, we focused our studies on female mice and did not test males for differences in sex effects. Thus, the female-specific conclusions we can draw from the data are limited without further studies. Future studies should assess the contribution of RGS14 to cocaine-taking behavior (e.g. self-administration) and include region- and cell type-specific manipulations to disentangle how RGS14 modulates motor and emotional circuits.

### 4.6. Conclusions

RGS14 suppresses synaptic plasticity in hippocampal area CA2 and hippocampal-based learning and memory. Our study shows that RGS14 is strongly expressed in discrete subnuclei of the NAc and extended amygdala, is found in D1R- and D2R-containing cells in the NAc, and is upregulated in the NAc in WT mice with a history of chronic cocaine. Behaviorally, loss of RGS14 increases cocaine-induced locomotor sensitization, CPP, and conditioned locomotor hyperactivity. Together, these findings point to a role for RGS14 as a natural barrier to plasticity in emotional and motivational brain circuits, and as a potentially protective factor against harmful cocaine-induced neuroadaptations leading to addiction.

## CREDIT AUTHORSHIP CONTRIBUTION STATEMENT: Sara N. Bramlett

Conceptualization, Methodology, Software, Formal analysis, Investigation, Writing-original draft, Visualization; **Stephanie L. Foster:** Conceptualization, Methodology, Formal analysis, Investigation; **David Weinshenker:** Conceptualization, Methodology, Resources, Writing-review & editing, Supervision, Funding acquisition; **John R. Hepler:** Conceptualization, Methodology, Resources, Writing-review & editing, Supervision, Funding acquisition.

## DATA AVAILABILITY

The raw data that support the findings of this study are available from the corresponding author, S.N.B., upon reasonable request.

## ACKNOWLEDGEMENTS

The authors would like to thank the Hess lab for their generous donation of *Drd1-tdTomato* and *Drd2-EGFP* tissue, Drs. Nicholas Harbin and Daniel Lustberg for helpful feedback and discussions regarding data and experimental planning, Dr. Daniel Lustberg for training in experimental techniques and assistance with materials, and Shana Fitzmaurice for assistance with running behavioral experiments.

## FUNDING AND DISCLOSURE

Research reported in this publication was supported in part by the Emory University Integrated Cellular Imaging Core of the Winship Cancer Institute of Emory University and NIH/NCI under award number, 2P30CA138292-04, (RRID:SCR_023534). This work was supported by the National Institutes of Health R01NS037112 (to J. R. H.), R01GM140632-A1 (to J. R. H and Peter A. Freidman). The authors have no conflicts of interest to declare.

## Notes

### Competing Interest Statement

The authors have declared no competing interest.

