## Supplemental Tables S1-S3 for "Endogenous Regulator of G protein Signaling 14 (RGS14) suppresses cocaine-induced emotionally motivated behaviors in female mice"

**Table S1. Locomotor sensitization statistics: induction.** Carat (^) indicates Holm correction.

| Fig | Statistical test | Day(s) | Factors | Effect or Groups | df | Statistic | Value | Pval | Sig |
|---|---|---|---|---|---|---|---|---|---|
| 4B | Linear Mixed-Effects models | Induction | Geno, Trtmt, Day | Geno | 1, 51.33 | F | 0.90 | .348 | |
| | | | | Trtmt | 1, 51.33 | F | 4.99 | .030 | * |
| | | | | Day | 1, 188 | F | 4.86 | .029 | * |
| | | | | Geno × Trtmt | 1, 51.33 | F | 3.43 | .070 | † |
| | | | | Geno × Day | 1, 188 | F | 6.78 | .010 | ** |
| | | | | Trtmt × Day | 1, 188 | F | 36.95 | < .001 | *** |
| | | | | Geno × Trtmt × Day | 1, 188 | F | 4.64 | .032 | * |
| 4C | Paired pairwise t-tests^ | Ind1 vs. Ind5 | | WT+Sal | 11 | t | 4.15 | .004 | ** |
| | | | | RGS14KO+Sal | 11 | t | 4.33 | .003 | ** |
| | | | | WT+CC | 11 | t | -1.63 | .131 | |
| | | | | RGS14KO+CC | 11 | t | -2.54 | .028 | * |
| 4D | Linear Mixed-Effects models | Induction | Trtmt, Day (WT) | Trtmt | 1, 29.99 | F | 0.21 | > .999 | |
| | | | | Day | 1, 94 | F | 0.12 | > .999 | |
| | | | | Trtmt × Day | 1, 94 | F | 11.56 | < .001 | *** |
| 4E | | | Trtmt, Day (RGS14-KO) | Trtmt | 1, 24.88 | F | 5.03 | .068 | † |
| | | | | Day | 1, 94 | F | 8.67 | .008 | ** |
| | | | | Trtmt × Day | 1, 94 | F | 25.41 | < .001 | *** |
| 4F | | | Geno, Day (Sal) | Geno | 1, 29.43 | F | 2.34 | .274 | |
| | | | | Day | 1, 94 | F | 23.29 | < .001 | *** |
| | | | | Geno × Day | 1, 94 | F | 0.31 | > .999 | |
| 4G | | | Geno, Day (Coc) | Geno | 1, 25.34 | F | 2.14 | .310 | |
| | | | | Day | 1, 94 | F | 20.45 | < .001 | *** |
| | | | | Geno × Day | 1, 94 | F | 6.75 | .022 | * |
| 4D | Pairwise t-tests^ | Ind1 | | WT+Coc vs. WT+Sal | 38.1 | t | 0.07 | .947 | |
| | | Ind2 | | | 38.1 | t | 1.02 | .633 | |
| | | Ind3 | | | 38.1 | t | 2.34 | .094 | † |
| | | Ind4 | | | 38.1 | t | 2.45 | .094 | † |
| | | Ind5 | | | 38.1 | t | 2.39 | .094 | † |
| 4E | | Ind1 | | RGS14KO+Coc vs. RGS14KO+Sal | 27.8 | t | 1.79 | .085 | † |
| | | Ind2 | | | 27.8 | t | 3.03 | .010 | * |
| | | Ind3 | | | 27.8 | t | 3.88 | .002 | ** |
| | | Ind4 | | | 27.8 | t | 3.80 | .002 | ** |
| | | Ind5 | | | 27.8 | t | 4.41 | < .001 | *** |
| 4G | | Ind1 | | RGS14KO+Coc vs. WT+Coc | 28.7 | t | 1.21 | .236 | |
| | | Ind2 | | | 28.7 | t | 1.95 | .144 | |
| | | Ind3 | | | 28.7 | t | 2.37 | .100 | † |
| | | Ind4 | | | 28.7 | t | 2.07 | .144 | |
| | | Ind5 | | | 28.7 | t | 2.80 | .045 | * |

**Table S2. Locomotor sensitization statistics: cocaine challenge.** Carat (^) indicates Holm correction.

| Fig | Statistical test | Day(s) | Factors | Effect or Groups | df | Statistic | Value | Pval | Sig |
|---|---|---|---|---|---|---|---|---|---|
| 4B | Two-Way ANOVA | Chall | Geno, Trtmt | Geno | 1, 44 | F | 4.98 | .031 | * |
| | | | | Trtmt | 1, 44 | F | 5.38 | .025 | * |
| | | | | Geno × Trtmt | 1, 44 | F | 4.63 | .037 | * |
| 4D | Tukey HSD test | Chall | | WT+Coc vs. WT+Sal | 44 | t | 0.71 | .894 | |
| 4E | | | | RGS14KO+Coc vs. RGS14KO+Sal | 44 | t | 3.41 | .008 | ** |
| 4F | | | | RGS14KO+Sal vs. WT+Sal | 44 | t | 0.64 | .918 | |
| 4G | | | | RGS14KO+Coc vs. WT+Coc | 44 | t | 3.34 | .009 | ** |
| 4H | Linear Mixed-Effects models | Chall | Geno, Trtmt, Time | Geno | 1, 418.62 | F | 14.35 | < .001 | *** |
| | | | | Trtmt | 1, 418.62 | F | 55.34 | < .001 | *** |
| | | | | Time | 1, 513 | F | 496.2 | < .001 | *** |
| | | | | Geno × Trtmt | 1, 418.62 | F | 24.67 | < .001 | *** |
| | | | | Geno × Time | 1, 513 | F | 8.65 | .003 | ** |
| | | | | Trtmt × Time | 1, 513 | F | 55.52 | < .001 | *** |
| | | | | Geno × Trtmt × Time | 1, 513 | F | 5.24 | < .001 | *** |
| 4I | Linear Mixed-Effects models | Chall (Habituation) | Trtmt, Time (WT) | Trtmt | 1, 33.33 | F | 1.07 | .309 | |
| | | | | Time | 1, 389 | F | 435.97 | < .001 | *** |
| | | | | Trtmt × Time | 1, 389 | F | 0.02 | .879 | |
| 4J | | | Trtmt, Time (RGS14-KO) | Trtmt | 1, 27.09 | F | 0.01 | .927 | |
| | | | | Time | 1, 406 | F | 413.84 | < .001 | *** |
| | | | | Trtmt × Time | 1, 406 | F | 3.54 | .061 | † |
| 4K | | | Geno, Time (Sal) | Geno | 1, 28.92 | F | 0.07 | .787 | |
| | | | | Time | 1, 389 | F | 438.79 | < .001 | *** |
| | | | | Geno × Time | 1, 389 | F | 0.88 | .350 | |
| 4L | | | Geno, Time (Coc) | Geno | 1, 28.22 | F | 0.41 | .530 | |
| | | | | Time | 1, 406 | F | 408.08 | < .001 | *** |
| | | | | Geno × Time | 1, 406 | F | 1.25 | .264 | |
| 4I | | Chall (Post-Coc) | Trtmt, Time (WT) | Trtmt | 1, 256.49 | F | 17.95 | < .001 | *** |
| | | | | Time | 1, 240 | F | 233.21 | < .001 | *** |
| | | | | Trtmt × Time | 1, 240 | F | 17.19 | < .001 | *** |
| 4J | | | Trtmt, Time (RGS14-KO) | Trtmt | 1, 184.89 | F | 61.05 | < .001 | *** |
| | | | | Time | 1, 262 | F | 267.08 | < .001 | *** |
| | | | | Trtmt × Time | 1, 262 | F | 61.43 | < .001 | *** |
| 4K | | | Geno, Time (Sal) | Geno | 1, 170.03 | F | 0.53 | .469 | |
| | | | | Time | 1, 240 | F | 132.30 | < .001 | *** |
| | | | | Geno × Time | 1, 240 | F | 0.04 | .846 | |
| 4L | | | Geno, Time (Coc) | Geno | 1, 228.52 | F | 32.60 | < .001 | *** |
| | | | | Time | 1, 262 | F | 365.82 | < .001 | *** |
| | | | | Geno × Time | 1, 262 | F | 23.81 | < .001 | *** |
| 4I | Pairwise t-tests^ | Chall | WT+Coc vs. WT+Sal | t = 95 min | 81.1 | t | 3.33 | .015 | * |
| | | | | t = 100 min | 81.1 | t | 3.17 | .023 | * |
| | | | | t = 105 min | 81.1 | t | 1.14 | > .999 | |
| | | | | t = 110 min | 81.1 | t | 1.47 | > .999 | |
| | | | | t = 115 min | 81.1 | t | 0.45 | > .999 | |
| | | | | t = 120 min | 81.1 | t | -0.92 | > .999 | |
| | | | | t = 125 min | 81.1 | t | -0.02 | > .999 | |
| | | | | t = 130 min | 81.1 | t | -0.02 | > .999 | |
| | | | | t = 135 min | 81.1 | t | -0.15 | > .999 | |
| | | | | t = 140 min | 81.1 | t | -0.07 | > .999 | |
| | | | | t = 145 min | 81.1 | t | 0.46 | > .999 | |
| | | | | t = 150 min | 81.1 | t | 1.17 | > .999 | |
| 4J | | | RGS14KO+Coc vs. RGS14KO+Sal | t = 95 min | 49.2 | t | 5.68 | < .001 | *** |
| | | | | t = 100 min | 49.2 | t | 4.25 | .001 | ** |
| | | | | t = 105 min | 49.2 | t | 2.64 | .099 | † |
| | | | | t = 110 min | 49.2 | t | 2.82 | .070 | † |
| | | | | t = 115 min | 49.2 | t | 1.54 | > .999 | |
| | | | | t = 120 min | 49.2 | t | 1.37 | > .999 | |
| | | | | t = 125 min | 49.2 | t | 0.42 | > .999 | |
| | | | | t = 130 min | 49.2 | t | 0.64 | > .999 | |
| | | | | t = 135 min | 49.2 | t | 0.37 | > .999 | |
| | | | | t = 140 min | 49.2 | t | 0.25 | > .999 | |
| | | | | t = 145 min | 49.2 | t | 0.12 | > .999 | |
| | | | | t = 150 min | 49.2 | t | 1.01 | > .999 | |
| 4L | | | RGS14KO+Coc vs. WT+Coc | t = 95 min | 57.1 | t | 4.31 | < .001 | *** |
| | | | | t = 100 min | 57.1 | t | 4.08 | .002 | ** |
| | | | | t = 105 min | 57.1 | t | 3.26 | .019 | * |
| | | | | t = 110 min | 57.1 | t | 2.63 | .076 | † |
| | | | | t = 115 min | 57.1 | t | 2.99 | .037 | * |
| | | | | t = 120 min | 57.1 | t | 2.89 | .044 | * |
| | | | | t = 125 min | 57.1 | t | 1.42 | .578 | |
| | | | | t = 130 min | 57.1 | t | 1.58 | .578 | |
| | | | | t = 135 min | 57.1 | t | 1.60 | .578 | |
| | | | | t = 140 min | 57.1 | t | 0.93 | .712 | |
| | | | | t = 145 min | 57.1 | t | 0.69 | .712 | |
| | | | | t = 150 min | 57.1 | t | 1.70 | .571 | |

### Table S3. Conditioned place preference statistics. Carat (^) indicates Bonferroni correction.

| Fig | Statistical test | Day(s) | Outcome measure | Factors (ANOVA) | Effect or Groups | df | Statistic | Value | Pval | Sig |
|---|---|---|---|---|---|---|---|---|---|---|
| 5B | Three-way ANOVA | | Time on Coc side (ΔPre-Test) | Geno, Trtmt, Stage | Geno | 1, 28 | F | 2.99 | .095 | † |
| | | | | | Trtmt | 1, 28 | F | 59.24 | < .001 | *** |
| | | | | | Stage | 1.54, 43.24 | F | 11.10 | < .001 | *** |
| | | | | | Geno × Trtmt | 1, 28 | F | 3.08 | .090 | † |
| | | | | | Geno × Stage | 1.54, 43.24 | F | 1.00 | .358 | |
| | | | | | Trtmt × Stage | 1.54, 43.24 | F | 10.78 | < .001 | *** |
| | | | | | Geno × Trtmt × Stage | 1.54, 43.24 | F | 1.02 | .353 | |
| | Two-way ANOVA | Post1 | Time on Coc side (ΔPre-Test) | Geno, Trtmt | Geno | 1, 28 | F | 1.84 | .186 | |
| | | | | | Trtmt | 1, 28 | F | 63.88 | < .001 | *** |
| | | | | | Geno × Trtmt | 1, 28 | F | 4.57 | .041 | * |
| | | Post2 | | Geno, Trtmt | Geno | 1, 28 | F | 5.16 | .031 | * |
| | | | | | Trtmt | 1, 28 | F | 74.40 | < .001 | *** |
| | | | | | Geno × Trtmt | 1, 28 | F | 3.26 | .082 | † |
| | | Post3 | | Geno, Trtmt | Geno | 1, 28 | F | 1.22 | .278 | |
| | | | | | Trtmt | 1, 28 | F | 20.45 | < .001 | *** |
| | | | | | Geno × Trtmt | 1, 28 | F | 0.81 | .377 | |
| | Tukey HSD test^ | Post1 | Time on Coc side (ΔPre-Test) | | KO+Sal vs. WT+Sal | 28 | t | -0.55 | .945 | |
| | | | | | WT+Coc vs. WT+Sal | 28 | t | 4.14 | .002 | ** |
| | | | | | KO+Coc vs. KO+Sal | 28 | t | 7.16 | < .001 | *** |
| | | | | | KO+Coc vs. WT+Coc | 28 | t | 2.47 | .087 | † |
| | | Post2 | | | KO+Sal vs. WT+Sal | 28 | t | 0.33 | .988 | |
| | | | | | WT+Coc vs. WT+Sal | 28 | t | 4.82 | < .001 | *** |
| | | | | | KO+Coc vs. KO+Sal | 28 | t | 7.38 | < .001 | *** |
| | | | | | KO+Coc vs. WT+Coc | 28 | t | 2.88 | .036 | * |
| | | Post3 | | | KO+Sal vs. WT+Sal | 28 | t | 0.15 | .999 | |
| | | | | | WT+Coc vs. WT+Sal | 28 | t | 2.56 | .072 | † |
| | | | | | KO+Coc vs. KO+Sal | 28 | t | 3.83 | .004 | ** |
| | | | | | KO+Coc vs. WT+Coc | 28 | t | 1.42 | .499 | |
| 5C | Three-way ANOVA | | Dist on Coc side (ΔPre-Test) (ranked) | Geno, Trtmt, Stage | Geno | 1, 28 | F | 2.89 | .100 | |
| | | | | | Trtmt | 1, 28 | F | 13.11 | .001 | ** |
| | | | | | Stage | 1.67, 46.86 | F | 0.00 | > .999 | |
| | | | | | Geno × Trtmt | 1, 28 | F | 1.64 | .211 | |
| | | | | | Geno × Stage | 1.67, 46.86 | F | 0.89 | .402 | |
| | | | | | Trtmt × Stage | 1.67, 46.86 | F | 3.17 | .060 | † |
| | | | | | Geno × Trtmt × Stage | 1.67, 46.86 | F | 0.45 | .604 | |
| | Two-way ANOVA | Post1 | Dist on Coc side (ΔPre-Test) (ranked) | Geno, Trtmt | Geno | 1, 28 | F | 5.05 | .033 | * |
| | | | | | Trtmt | 1, 28 | F | 29.45 | < .001 | *** |
| | | | | | Geno × Trtmt | 1, 28 | F | 3.08 | .090 | † |
| | | Post2 | | Geno, Trtmt | Geno | 1, 28 | F | 3.04 | .092 | † |
| | | | | | Trtmt | 1, 28 | F | 6.96 | .013 | * |
| | | | | | Geno × Trtmt | 1, 28 | F | 1.52 | .227 | |
| | | Post3 | | Geno, Trtmt | Geno | 1, 28 | F | 0.55 | .464 | |
| | | | | | Trtmt | 1, 28 | F | 4.46 | .044 | * |
| | | | | | Geno × Trtmt | 1, 28 | F | 0.34 | .562 | |
| | Tukey HSD test^ | Post1 | Dist on Coc side (ΔPre-Test) (ranked) | | KO+Sal vs. WT+Sal | 28 | t | 0.35 | .985 | |
| | | | | | WT+Coc vs. WT+Sal | 28 | t | 2.60 | .067 | † |
| | | | | | KO+Coc vs. KO+Sal | 28 | t | 5.08 | < .001 | *** |
| | | | | | KO+Coc vs. WT+Coc | 28 | t | 2.83 | .040 | * |
| | | Post2 | | | KO+Sal vs. WT+Sal | 28 | t | 0.36 | .984 | |
| | | | | | WT+Coc vs. WT+Sal | 28 | t | 0.99 | .755 | |
| | | | | | KO+Coc vs. KO+Sal | 28 | t | 2.74 | .049 | * |
| | | | | | KO+Coc vs. WT+Coc | 28 | t | 2.11 | .176 | |
| | | Post3 | | | KO+Sal vs. WT+Sal | 28 | t | 0.11 | > .999 | |
| | | | | | WT+Coc vs. WT+Sal | 28 | t | 1.08 | .706 | |
| | | | | | KO+Coc vs. KO+Sal | 28 | t | 1.91 | .248 | |
| | | | | | KO+Coc vs. WT+Coc | 28 | t | 0.94 | .784 | |
| N/A | Three-way ANOVA | | Total dist (ΔPre-Test) (ranked) | Geno, Trtmt, Stage | Geno | 1, 28 | F | 1.94 | .175 | |
| | | | | | Trtmt | 1, 28 | F | 0.09 | .762 | |
| | | | | | Stage | 1.73, 48.53 | F | 0.00 | > .999 | |
| | | | | | Geno × Trtmt | 1, 28 | F | 0.47 | .497 | |
| | | | | | Geno × Stage | 1.73, 48.53 | F | 5.33 | .011 | * |
| | | | | | Trtmt × Stage | 1.73, 48.53 | F | 5.54 | .009 | ** |
| | | | | | Geno × Trtmt × Stage | 1.73, 48.53 | F | 0.87 | .411 | |
| | Two-way ANOVA | Post1 | Total dist (ΔPre-Test) (ranked) | Geno, Trtmt | Geno | 1, 28 | F | 4.70 | .039 | * |
| | | | | | Trtmt | 1, 28 | F | 2.09 | .159 | |
| | | | | | Geno × Trtmt | 1, 28 | F | 0.41 | .526 | |
| | | Post2 | | Geno, Trtmt | Geno | 1, 28 | F | 4.38 | .046 | * |
| | | | | | Trtmt | 1, 28 | F | 2.83 | .104 | |
| | | | | | Geno × Trtmt | 1, 28 | F | 1.51 | .229 | |
| | | Post3 | | Geno, Trtmt | Geno | 1, 28 | F | 0.26 | .617 | |
| | | | | | Trtmt | 1, 28 | F | 0.29 | .592 | |
| | | | | | Geno × Trtmt | 1, 28 | F | 0.00 | .971 | |
| | Tukey HSD test^ | Post1 | Total dist (ΔPre-Test) (ranked) | | KO+Sal vs. WT+Sal | 28 | t | 1.08 | .705 | |
| | | | | | WT+Coc vs. WT+Sal | 28 | t | 0.57 | .941 | |
| | | | | | KO+Coc vs. KO+Sal | 28 | t | 1.48 | .464 | |
| | | | | | KO+Coc vs. WT+Coc | 28 | t | 1.99 | .217 | |
| | | Post2 | | | KO+Sal vs. WT+Sal | 28 | t | 0.61 | .928 | |
| | | | | | WT+Coc vs. WT+Sal | 28 | t | -2.06 | .191 | |
| | | | | | KO+Coc vs. KO+Sal | 28 | t | -0.32 | .989 | |
| | | | | | KO+Coc vs. WT+Coc | 28 | t | 2.35 | .111 | |
| | | Post3 | | | KO+Sal vs. WT+Sal | 28 | t | -0.33 | .987 | |
| | | | | | WT+Coc vs. WT+Sal | 28 | t | -0.36 | .984 | |
| | | | | | KO+Coc vs. KO+Sal | 28 | t | -0.41 | .976 | |
| | | | | | KO+Coc vs. WT+Coc | 28 | t | -0.38 | .980 | |
| 5D | Pearson's correlation | Post1 | ΔTime vs. ΔDist correlation | | WT | 6 | t | 2.32 | .059 | † |
| | | | | | | | R | 0.69 | | |
| | | | | | KO | 6 | t | 0.83 | .439 | |
| | | | | | | | R | 0.32 | | |
| 5E | Welch Two Sample t-test | Pre-Test | Time on white-floored side | | KO vs. WT | 28.82 | t | -3.79 | < .001 | *** |
| 5F | Welch Two Sample t-test | Post1 | Time on white-floored side | (Coc group) | KO vs. WT | 9.28 | t | 1.33 | .214 | |
